# Terrestrial force production by the limbs of a semi-aquatic salamander provides insight into the evolution of terrestrial locomotor mechanics

**DOI:** 10.1101/2021.05.01.442256

**Authors:** Sandy M. Kawano, Richard W. Blob

## Abstract

Amphibious fishes and salamanders are valuable functional analogs for vertebrates that spanned the water-to-land transition. However, investigations of walking mechanics have focused on terrestrial salamanders and, thus, may better reflect the capabilities of stem tetrapods that were already terrestrial. The earliest tetrapods were aquatic, so salamanders that are not primarily terrestrial may yield more appropriate data for modelling the incipient stages of terrestrial locomotion. In the present study, locomotor biomechanics were quantified from semi-aquatic *Pleurodeles waltl*, a salamander that spends most of its adult life in water, and then compared to a primarily terrestrial salamander (*Ambystoma tigrinum*) and semi-aquatic fish (*Periophthalmus barbarus*) to evaluate whether walking mechanics show greater similarity between species with ecological versus phylogenetic similarities. Ground reaction forces (GRFs) from individual limbs or fins indicated that the pectoral appendages of each taxon had distinct patterns of force production, but hind limb forces were comparable between the salamanders. The rate of force development (‘yank’) was sometimes slower in *P. waltl* but generally comparable between the three species. Finally, medial inclination of the GRF in *P. waltl* was intermediate between semi-aquatic fish and terrestrial salamanders, potentially elevating bone stresses among more aquatic taxa as they move on land. These data provide a framework for modelling stem tetrapods using an earlier stage of quadrupedal locomotion that was powered primarily by the hind limbs (i.e., “rear-wheel drive”), and reveal mechanisms for appendages to generate propulsion in three locomotor strategies that are presumed to have occurred across the water-to-land transition in vertebrate evolution.

**Summary statement:** Semi-aquatic salamanders had limb mechanics that were intermediate in magnitude yet steadier than the appendages of terrestrial salamanders and semi-aquatic fish, providing a framework to model semi-aquatic early tetrapods.

## INTRODUCTION

The evolutionary invasion of land was a seminal event in vertebrate history that has received intense study, with integrative approaches filling major gaps in our understanding of how this transition transpired (Ashley-Ross et al. 2013). Skeletal remains and trackways in the fossil record can provide some of the most direct evidence about such historical events, yet also present challenges given the limits to which fossils preserve behavior and functional performance (see (Gatesy et al., 1999; Maidment et al., 2014); and references therein). Consequently, extant amphibious fishes, amphibians, and reptiles have been used as functional models to infer the biology of extinct tetrapodomorphs (tetrapods and tetrapod-like sarcopterygian fishes) (Nyakatura et al., 2019, 2014; Pierce et al., 2013), with extant taxa representing alternative strategies for invading land and potentially serving as models for different functional stages as vertebrates became terrestrial. Investigations of extant taxa exhibiting morphological and/or behavioral traits that are consistent with those of extinct tetrapodomorphs offer particularly intriguing opportunities to evaluate how tetrapods were able to leave the water’s edge (Pierce et al., 2013).

In considerations of locomotor performance during the invasion of land, salamanders are often used as functional analogues for early tetrapods since they move between water and land (Karakasiliotis et al., 2012), and exhibit a relatively generalized tetrapod *Bauplan* that has not changed substantially for at least 150 million years (Gao and Shubin, 2001) and possibly as long as 230 million years (Schoch et al., 2020). Previous studies used living salamanders to gain insights into the biomechanics and muscle physiology of walking underwater (Ashley-Ross et al., 2009; Azizi and Horton, 2004; Deban and Schilling, 2009; Frolich and Biewener, 1992) and on land (Ashley-Ross et al., 2009; Brand, 1996; Deban and Schilling, 2009; Delvolvé et al., 1997; Frolich and Biewener, 1992; Kawano and Blob, 2013; Pierce et al., 2020; Sheffield and Blob, 2011), transitioning between water and land (Ashley-Ross and Bechtel, 2004), and assessing how bone histology relates to ecological habits (Canoville and Laurin, 2009; Laurin et al., 2004). Given the greater effect of gravitational loads on the musculoskeletal system in terrestrial environments (Young et al., 2017), one of the most fundamental requirements for moving on land is the ability to support body weight for maintaining posture and physical forces for locomotion. Evaluations of the weight-bearing capabilities in the limbs of early tetrapods have been approached through measurements of ground reaction forces (GRFs) experienced by the tiger salamander, *Ambystoma tigrinum*, which demonstrated that the hind limbs supported a similar proportion of body weight as the forelimbs but had a greater contribution to acceleration (Kawano and Blob, 2013). To what extent might these results relate to the locomotor function of early tetrapods? Fossil evidence suggests that some of the first tetrapods, such as *Acanthostega*, were still aquatic (Coates, 1996), and other early tetrapods, such as *Ichthyostega*, may have had only limited terrestrial capabilities (Pierce et al., 2012). In contrast, *A. tigrinum* are found in various terrestrial habitats, ranging from conifer forests to deserts, and adults rarely spend extended durations in water for reasons other than reproduction (Petranka, 1998). As such, they may not provide an ideal model for the initial invaders of land, in which terrestrial capacity may not have been fully acquired. How might limb function differ for a species that spends most of its time in water?

Because salamanders have a diverse range of habitat preferences and life histories (Wake, 2009), they provide an opportunity to model different evolutionary stages in the adoption of terrestrial habits. Examinations of amphibious species with greater aquatic tendencies could yield substantial insights into the limb function of earlier stem tetrapods with digit-bearing limbs. In this study, we compared GRFs of individual limbs produced by semi-aquatic salamanders (Spanish ribbed newts, *Pleurodeles waltl* Michahelles 1830; see Electronic Supplementary Materials for details), to published data on terrestrial tiger salamanders, *Ambystoma tigrinum* Green 1825, and semi-aquatic African mudskippers, *Periophthalmus barbarus* (Linnaeus 1766) (Kawano and Blob, 2013). We also calculated new measures of yank, the rate of force development (*sensu* Lin et al., 2019), to evaluate how the rate of force production might differ between these species. Quantifying the dynamics of force-time measures contributes valuable information to explain kinematic and muscle activation patterns (Lin et al., 2019). Our aim in these comparisons was to examine extant taxa that model important stages during the transition to land to quantify the functional differences between fins and limbs for terrestrial locomotion.

During aquatic locomotion, dominant force production by the rear appendage (“rear-wheel drive”) may have appeared as early as in sarcopterygian fishes (King et al., 2011). However, terrestrial propulsion in stem tetrapods may have been dominated by the forelimb (‘front-wheel drive’), with the transition to hind limb dominance (‘rear-wheel drive’) only occurring among lineages that diverged later, as the hind limbs assumed a more important locomotor role (Boisvert, 2005). The ‘front-wheel drive’ strategy proposed for the sarcopterygian fish *Panderichthys* (Boisvert, 2005) and the early tetrapod *Ichthyostega* (Pierce et al., 2012) might be appropriately modeled by the locomotor behaviors of extant mudskipper fishes (McInroe et al., 2016), which use synchronous movements of the pectoral fins to ‘crutch’ over land while keeping the body axis relatively straight (Kawano and Blob, 2013; Pace and Gibb, 2014). On the other hand, the ‘rear-wheel drive’ of more crownward early tetrapods might be better modeled using terrestrial salamanders, such as *A. tigrinum* (Pierce et al., 2013). Our functional models also show distinct GRF patterns between these groups, with those produced by the pectoral fins of mudskippers directed more medially (~17°) than those produced by the limbs of terrestrial salamanders, or almost any other terrestrial tetrapod (generally less than 11°). Such differences could have substantial implications for the stresses experienced by the limb bones by increasing moment arms due to bending, which could make bones more vulnerable to fracture (Kawano and Blob, 2013).

Our data from semi-aquatic *P. waltl* can give new insight into the functional changes that occurred during the early evolution of terrestrial locomotion. Simply by having limbs, locomotor force production by *P. waltl* may be similar to that of terrestrial salamanders such as adult *A. tigrinum*. However, the habitual use of the limbs for aquatic locomotion by adult *P. waltl* might lead to kinetic similarities with other semi-aquatic taxa, such as mudskipper fishes. These comparisons carry broader implications for generating hypotheses about the emergence of functional disparity between locomotor structures, such as whether changes in functional performance are coupled across major structural changes (e.g., transition from fin to limb) or through gradual steps related to loading regimes that are potentially decoupled from structural changes (e.g., the transition from aquatic to terrestrial habitats, regardless of which locomotor structure is used).

## MATERIALS AND METHODS

### Animals

Five adult *P. waltl* (body mass: 16.60 ± 0.40 g; snout-vent length: 0.083 ± 0.001 m; total length: 0.186 ± 0.003 m) were obtained from a commercial vendor. All values represent means ± 1 S.E. Animals were individually housed in glass aquaria that were aerated with sponge filters, kept on a 12h:12h light:dark cycle, and fed every one - two days on a diet of defrosted bloodworms and krill. Prior to testing, animals were starved for two days to reduce the effects of satiation on locomotor performance (Sass and Motta, 2002).

### Collection of three-dimensional (3D) ground reaction forces

Experimental procedures from a previous study on the GRFs of tiger salamanders and mudskipper fish (Kawano and Blob, 2013) were replicated in the present study (see Electronic Supplementary Materials) to obtain forelimb (*n* = 50) and hind limb (*n* = 49) GRFs from *P. waltl* (Fig. S1). Custom code to replicate these analyses is available through GitHub: https://github.com/MorphoFun/kraken. The focal taxa examined in the present study represent models for distinct functional stages during the evolution of terrestrial locomotion: front-wheel drive in a semi-aquatic vertebrate with limited capabilities of the pelvic appendages (semi-aquatic mudskipper fish), a semi-aquatic vertebrate with a generalized tetrapod *Bauplan* (semi-aquatic *P. waltl* salamander), and rear-wheel drive in a terrestrial vertebrate with a generalized tetrapod *Bauplan* (terrestrial *A. tigrinum* salamander). GRFs in the vertical, mediolateral, and anteroposterior directions were digitally filtered with a custom low-pass, zero phase second order Butterworth filter, and then interpolated to 101 points (0-100% of stance at 1% increments) using a cubic spline with the *signal* R package (Signal developers, 2013). The stance phase was defined to begin at the first video frame when the entire foot was flat against the ground, and to end at the frame penultimate to the beginning of the swing phase.

Interspecific and intraspecific comparisons were made across the GRFs of fins and limbs. First, GRFs from the forelimbs and hind limbs of *P. waltl* were compared to published data from the pectoral and pelvic appendages of mudskipper fish and tiger salamanders (Kawano and Blob, 2013) to assess whether limb kinetics in semi-aquatic salamanders are more similar to those of mudskipper fins or terrestrial salamander limbs. Second, GRFs were compared between the forelimbs and hind limbs of *P. waltl* to determine whether a model for a semi-aquatic early tetrapod was forelimb-driven or hind limb-driven. Third, GRF data from these three species were used to conduct new analyses of yank (*sensu* Lin et al., 2019) to quantify the rate of change in the GRFs produced by fins vs. limbs. Linear mixed effects models (LMMs) were used to compare magnitudes and angles of orientation of GRFs when forces were maximal (“peak net GRF”), and the range of yank values recorded between the species and appendages (see “Statistical analyses”).

### Statistical analyses

Linear mixed effects models with random intercepts via the *lme4* package (Bates et al., 2015) were used to compare the biomechanical variables of interest while accounting for the non-independence of multiple trials sampled per individual, and fit with Restricted Maximum Likelihood (REML) to produce unbiased estimates. LMMs were conducted to account for the non-independence caused by trials being nested within individuals and the unequal sample sizes across individuals. LMMs also have the advantage that they do not make assumptions about balanced designs, homogeneity of variance, etc., so the model assumptions only include random sampling and normality of the residuals (reviewed in (Smith, 2017). Evaluation of Quantile-Quantile (Q-Q) plots indicated that there were no major deviations from the reference line (Electronic Supplementary Material), suggesting that all variables reasonably met the assumption of normality. However, violations of the model assumptions would not have affected our analyses given our focus on fixed effects, which tend to be unbiased even if assumptions are violated (LeBeau et al., 2018). “Individual” was treated as a random effect, and “appendage type” (pectoral vs. pelvic) or “species” was treated as a fixed effect for intra- and interspecific comparisons, respectively.

Large sample sizes are typically required to produce accurate estimates of a random slopes model (reviewed in Harrison et al. 2018) so we assumed that the slopes were similar across the individuals within a species based on our smaller sample size. Assuming common slopes in a LMM can increase the potential for false positives and false negatives when calculating p-values for null hypothesis testing (reviewed in Harrison et al. 2018); however, the present study focused on effect sizes (i.e., the magnitude of the effects or variables) rather than the p-values, so these outcomes do not detract from our analyses. Given that one of the objectives of this study is to quantify the mean differences (i.e., point estimates) and range of variation (i.e., interval estimates) in GRF production between and within salamanders to parameterize computational models of musculoskeletal function, our emphasis on calculating effect sizes is appropriate (Cohen, 1990) due to the limited information that often results from traditional methods of null hypothesis testing (Cumming, 2014).

To calculate the point estimates (i.e., mean values) and interval estimates (i.e., standard errors) of the fixed effects, we used the emmeans::emmeans() function (Lenth, 2019). The coefficient of determination (percent of the variation explained by each model) was calculated as Nagakawa’s marginal R^2^ (R^2^_LMM(m)_; variation due to the fixed effects only) and conditional R^2^ (R^2^_LMM(c)_; variation due to the fixed and random effects) using the performance::r2_nagakawa() function (Lüdecke et al., 2019). Dynamic changes in force magnitudes and yank were plotted by pooling individual trials for a given species, but results from the LMMs are reported in the tables to account for repeated trials within individuals. All analyses were conducted in *R* version 3.6.1 (R Core Team, 2019) and *RStudio* version 1.2.1335 (RStudio Team, 2015).

## RESULTS

### Force production between the pectoral appendages of mudskippers and salamanders

The forelimbs of *P. waltl* exhibited kinetics that were generally intermediate between the pectoral fins of *P. barbarus* fish and the forelimbs of *A. tigrinum* salamanders (Fig. 1). Force production during terrestrial locomotion generally approximated a bell-shape curve that was skewed to the right in *P. barbarus* and *A. tigrinum*, but relatively broad and shallow in *P. waltl* for the net GRF and the vertical component of the GRF (Fig. 1a-b). The mediolateral component of the GRF was directed medially during stance for all three species, yet had a wider range of values in *P. barbarus* (Fig. 1c). For the anteroposterior component of the GRF, forelimb kinetics in *P. waltl* were more similar to terrestrial *A. tigrinum* salamanders than to semi-aquatic *P. barbarus* fish. The pectoral fin is the primary propulsor during crutching for *P. barbarus* and its GRF was oriented anteriorly for all of stance (indicating an acceleratory role), whereas the forelimbs of *A. tigrinum* and *P. waltl* showed primarily posteriorly directed GRFs (deceleratory role), with only one or two periods in which the GRFs became slightly anterior (Fig. 1d, f). The GRF was angled medially in the horizontal plane throughout stance for *P. barbarus* and *A. tigrinum*, and for almost all of stance for *P. waltl* except for a slight lateral orientation starting around 90% of stance (Fig. 1e). These findings illustrate how different aspects of force production by the forelimbs of *P. waltl* share similarities, yet also distinct differences (e.g., mediolateral GRFs), with those of semi-aquatic fins and terrestrial limbs.

**Figure 1.**
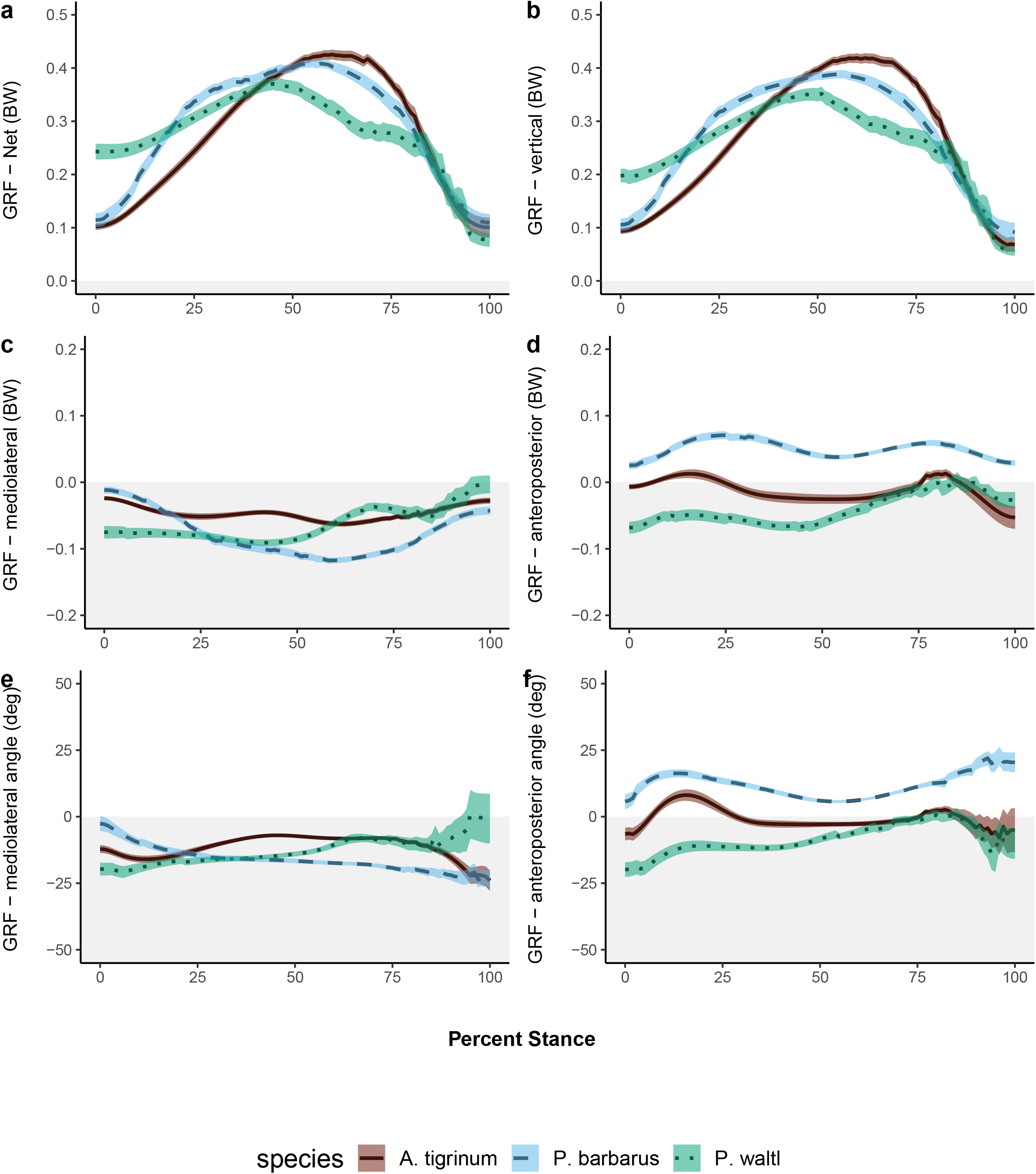
Profiles of GRF production in the pectoral appendages during stance. Individual trials were pooled within each species (N = 50 trials for each) and profiles are plotted as the mean ± s.e.m. for each percent of stance. Mediolateral angles were set relative to vertical (0°), so negative values indicate a medial direction of the GRF. Anteroposterior angles were set relative to vertical (0°), so negative values represent a posterior direction of the GRF. Although the general shapes of the curves were comparable across the species for a given parameter, pectoral appendages for *P. waltl* (dotted green lines) tended to support lower magnitudes of body weight, and have a more deceleratory role and intermediate mediolateral angle, compared to the semi-aquatic *P. barbarus* (dashed blue lines) and the primarily terrestrial *A. tigrinum* (solid brown lines).

GRF parameters at the peak net GRF also differed between the pectoral appendages for the three groups examined. The timing of the peak net GRF occurred at 60.4 ± 2.86 % of stance for *A. tigrinum*, 56.0 ± 2.86 % for *P. barbarus,* and 46.9 ± 2.93 % for *P. waltl* (mean ± SE; R^2^_LMM(m)_ = 0.226, R^2^_LMM(c)_ = 0.462). The magnitude of the peak net GRF was comparable between *P. barbarus* and *A. tigrinum* at approximately 44% and 46% of body weight, respectively, but was roughly 40% of body weight and had a wider range of values in *P. waltl* (Table 1, Fig. 2a). Similarly, the vertical component at the peak net GRF was comparable for *A. tigrinum* and *P. barbarus* yet lower and broader for *P. barbarus* (Fig. 2b). The mediolateral component of the peak net GRF was similar between the two salamanders (Fig. 2c), with values for *P. barbarus* being 1.5 – 2x more medial than the salamanders. The anteroposterior component was also more similar between the salamanders at the peak net GRF, with values being anterior (acceleratory) for *P. barbarus* but posterior (deceleratory) for both salamanders. The angle of the GRF was medial for all pectoral appendages at peak net GRF, but values for *P. waltl* were intermediate (~14°) between *P. barbarus* (~17°) and *A. tigrinum* (~9°). These comparisons further demonstrate that kinetics at the time of peak net GRF for *P. waltl* share similarities to semi-aquatic fins and terrestrial limbs, with some features intermediate between the other two species.

**Figure 2.**
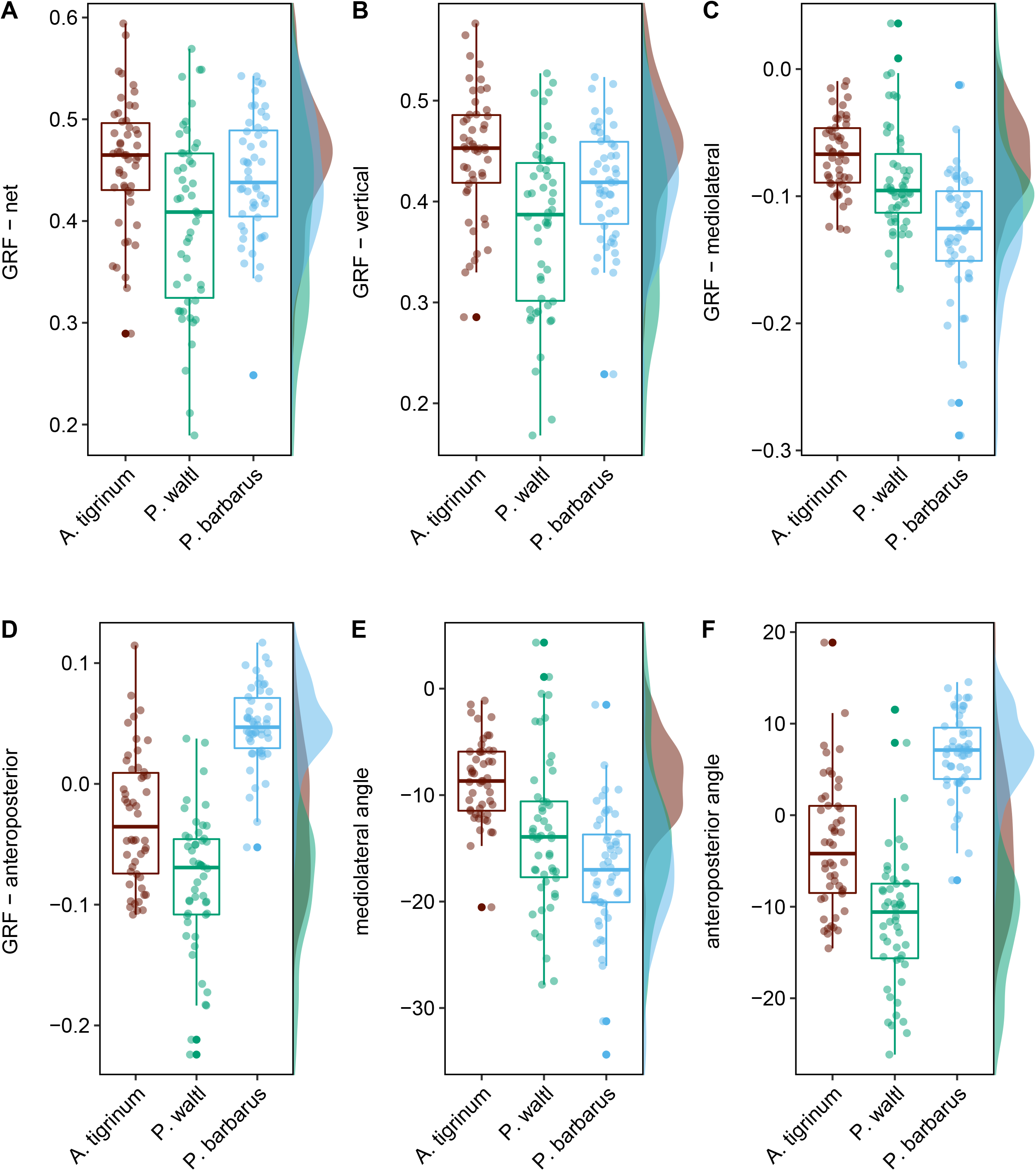
Parameters at the peak net GRF compared across the pectoral appendages. *P. waltl* (green) generally has lower magnitudes for the vertical component and net GRF as demonstrated by the histograms, yet also had a greater variability compared to the semi-aquatic fish *P. barbarus* (blue) and the primarily terrestrial salamander *A. tigrinum* (brown) as illustrated by the whisker-box plots. Sample sizes for data from the pectoral appendages: *P. barbarus* (n = 50), *P. waltl* (n = 50), and *A. tigrinum* (n = 50).

**Table 1.**
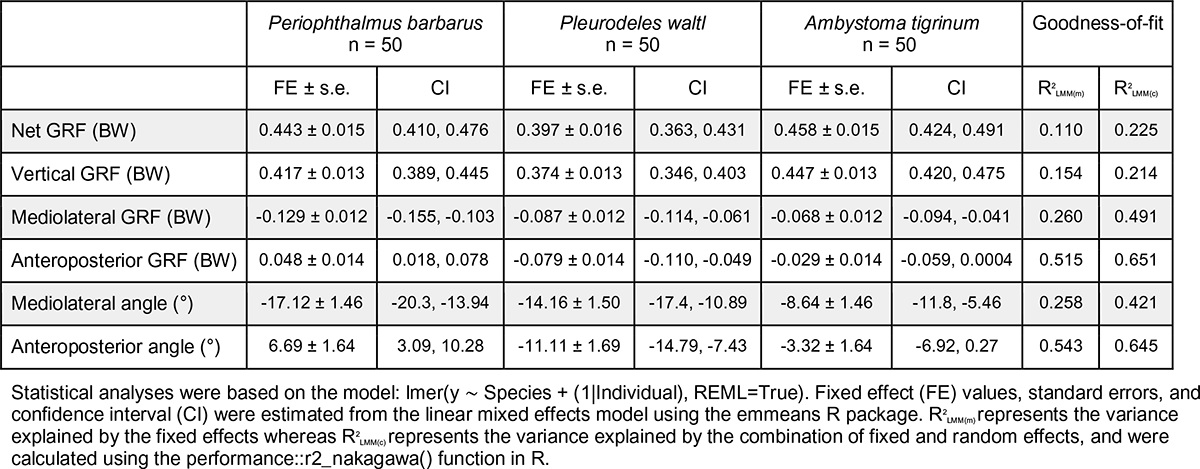
Comparison of locomotor variables between the pectoral appendages of mudskippers and salamanders at the peak net GRF

|  | *Periophthalmus barbarus*<br>n = 50 |  | *Pleurodeles waltl*<br>n = 50 |  | *Ambystoma tigrinum*<br>n = 50 |  | Goodness-of-fit |  |
| --- | --- | --- | --- | --- | --- | --- | --- | --- |
| | FE ± s.e. | CI | FE ± s.e. | CI | FE ± s.e. | CI | $R^2_{LMM(m)}$ | $R^2_{LMM(c)}$ |
| Net GRF (BW) | 0.443 ± 0.015 | 0.410, 0.476 | 0.397 ± 0.016 | 0.363, 0.431 | 0.458 ± 0.015 | 0.424, 0.491 | 0.110 | 0.225 |
| Vertical GRF (BW) | 0.417 ± 0.013 | 0.389, 0.445 | 0.374 ± 0.013 | 0.346, 0.403 | 0.447 ± 0.013 | 0.420, 0.475 | 0.154 | 0.214 |
| Mediolateral GRF (BW) | -0.129 ± 0.012 | -0.155, -0.103 | -0.087 ± 0.012 | -0.114, -0.061 | -0.068 ± 0.012 | -0.094, -0.041 | 0.260 | 0.491 |
| Anteroposterior GRF (BW) | 0.048 ± 0.014 | 0.018, 0.078 | -0.079 ± 0.014 | -0.110, -0.049 | -0.029 ± 0.014 | -0.059, 0.0004 | 0.515 | 0.651 |
| Mediolateral angle (°) | -17.12 ± 1.46 | -20.3, -13.94 | -14.16 ± 1.50 | -17.4, -10.89 | -8.64 ± 1.46 | -11.8, -5.46 | 0.258 | 0.421 |
| Anteroposterior angle (°) | 6.69 ± 1.64 | 3.09, 10.28 | -11.11 ± 1.69 | -14.79, -7.43 | -3.32 ± 1.64 | -6.92, 0.27 | 0.543 | 0.645 |

Statistical analyses were based on the model: lmer(y ~ Species + (1|Individual), REML=True). Fixed effect (FE) values, standard errors, and confidence interval (CI) were estimated from the linear mixed effects model using the emmeans R package. $R^2_{LMM(m)}$ represents the variance explained by the fixed effects whereas $R^2_{LMM(c)}$ represents the variance explained by the combination of fixed and random effects, and were calculated using the performance::r2\_nakagawa() function in R.

### Force production between salamander forelimbs and hind limbs

Kinetic profiles were similar in shape between the hind limbs of semi-aquatic and terrestrial salamanders (Fig. 3), although there were differences in the magnitudes of the curves. Semi-aquatic and terrestrial hind limbs had a peak net GRF occurring around 30% of stance; however, the net GRF and the vertical component at peak GRF for *P. waltl* were more similar to those of *P. waltl* forelimbs (at about 30% of body weight) than to the hind limbs of *A. tigrinum* (~50% of body weight) (Tables 1-2, Fig. 4a - b). The GRFs were directed medially for all limbs (~9-17°) but were 1.5x and 2x more medial in *P. waltl* than *A. tigrinum* for the forelimbs and hind limbs, respectively (Table 2, Fig. 4e). The forelimbs of both salamander species had a net deceleratory role (Table 1), whereas the hind limbs had a net acceleratory role (Table 2), with the hind limbs of *A. tigrinum* showing a mean value ~1.5× higher than those of *P. waltl* at the time of peak net GRF (Fig. 4d). However, the angle of the GRF in the anteroposterior direction was comparable between salamander hind limbs, inclined 16 – 17° anteriorly (Fig. 4f). These findings demonstrate that hind limb kinetics are not necessarily equivalent between salamander taxa with differing degrees of terrestriality, and that locomotor biomechanics differ between the forelimbs and hind limbs during terrestrial walking in both species.

**Figure 3.**
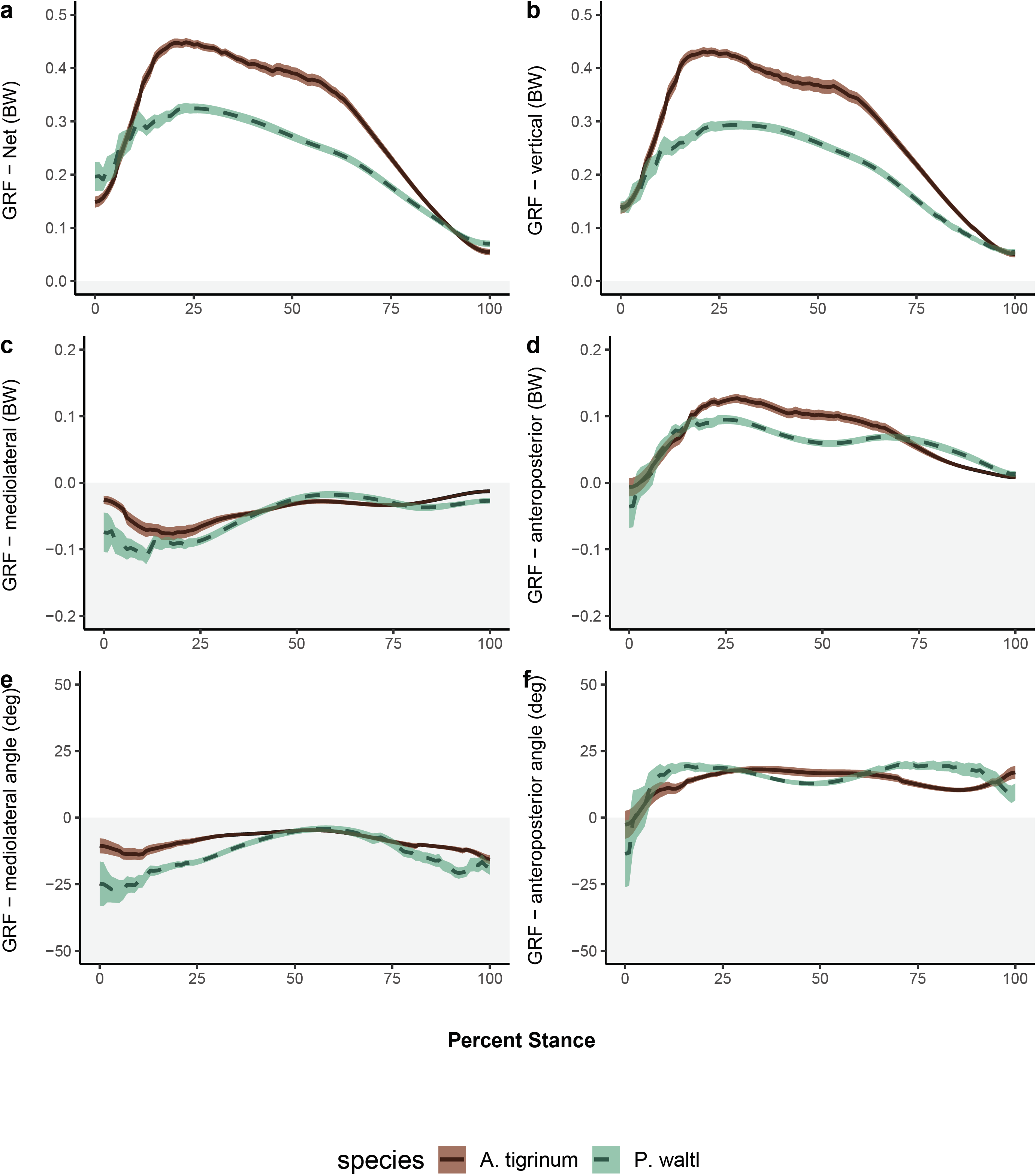
Profiles of GRF production in the pelvic appendages during stance. Individual trials were pooled within each species (N = 49 trials for each) and profiles are plotted as the mean ± s.e.m. for each percent of stance. Mediolateral angles were set relative to vertical (0°), so negative values indicate a medial direction of the GRF. Anteroposterior angles were set relative to vertical (0°), so negative values represent a posterior direction of the GRF. Although the general shapes of the curves were comparable across the species for a given parameter, *P. waltl* hind limbs (dashed green lines) tended to support lower magnitudes of body weight and had less of an acceleratory role compared to the primarily terrestrial salamander *A. tigrinum* (solid brown lines).

**Figure 4.**
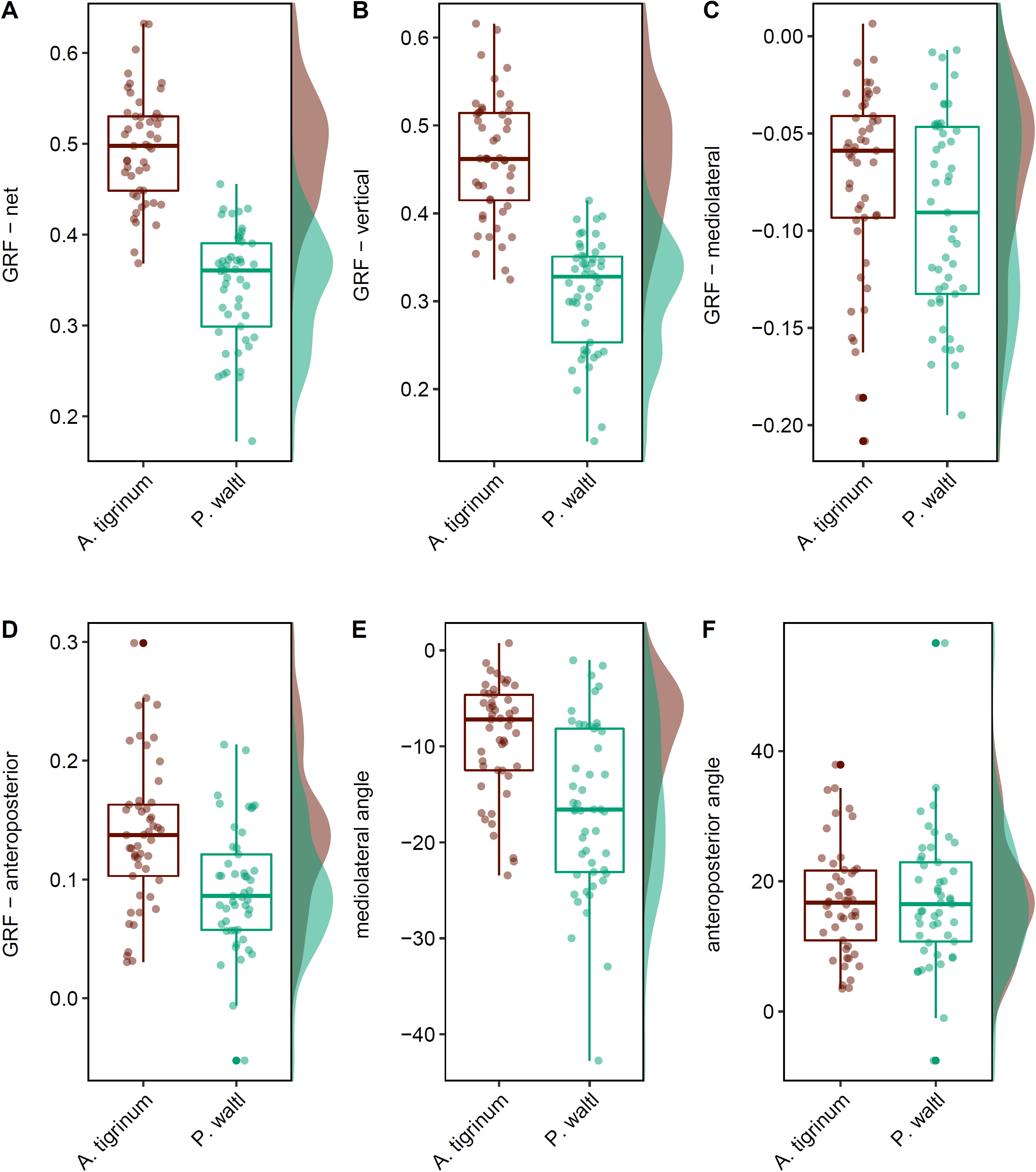
Parameters at the peak net GRF compared across the pelvic appendages. *P. waltl* hind limbs (green) generally had lower magnitudes for the vertical component and net GRF as demonstrated by the histograms, yet the angle of GRF orientation could become almost 2x more medial than for the primarily terrestrial salamander *A. tigrinum* (brown) as illustrated by the whisker-box plots. Sample sizes for the pelvic appendages: *P. waltl* (n = 49), and *A. tigrinum* (n = 49).

**Table 2.** Comparison of locomotor variables between the pelvic appendages of salamanders at the peak net GRF

|  | *Pleurodeles waltl*<br>n = 49 |  | *Ambystoma tigrinum*<br>n = 49 |  | Goodness-of-fit |  |
| --- | --- | --- | --- | --- | --- | --- |
| | FE ± s.e. | CI | FE ± s.e. | CI | R<sup>2</sup><sub>LMM(m)</sub> | R<sup>2</sup><sub>LMM(c)</sub> |
| Net GRF (BW) | 0.349 ± 0.017 | 0.309, 0.389 | 0.497 ± 0.017 | 0.457, 0.537 | 0.585 | 0.715 |
| Vertical GRF (BW) | 0.314 ± 0.021 | 0.266, 0.362 | 0.466 ± 0.021 | 0.419, 0.514 | 0.558 | 0.734 |
| Mediolateral GRF (BW) | -0.096 ± 0.014 | -0.129, -0.063 | -0.071 ± 0.014 | -0.104, -0.037 | 0.056 | 0.350 |
| Anteroposterior GRF (BW) | 0.092 ± 0.017 | 0.053, 0.132 | 0.132 ± 0.017 | 0.092, 0.172 | 0.102 | 0.418 |
| Mediolateral angle (°) | -16.97 ± 2.27 | -22.2, -11.7 | -8.73 ± 2.27 | -14.0, -3.5 | 0.215 | 0.483 |
| Anteroposterior angle (°) | 17.0 ± 2.71 | 10.7, 23.2 | 16.3 ± 2.70 | 10.0, 22.5 | 0.001 | 0.315 |

Statistical analyses were based on the model: lmer(y ~ Species + (1|Individual), REML=True). Fixed effect (FE) values, standard errors, and confidence interval (CI) were estimated from the linear mixed effects model using the emmeans R package. R<sup>2</sup><sub>LMM(m)</sub> represents the variance explained by the fixed effects whereas R<sup>2</sup><sub>LMM(c)</sub> represents the variance explained by the combination of fixed and random effects, and were calculated using the performance::r2\_nakagawa() function in R.

### Rate of change in GRF production

Yank values for the pectoral appendages tended to be similar between the three species, but GRFs produced by the forelimbs of *P. waltl* generally reached their peak more slowly (as indicated by smaller absolute magnitudes of yank) and had a narrower range compared to the semi-aquatic fish and terrestrial salamanders (Table 3). The shapes of the curves for yank differed between *P. waltl* and the other two species. For *P. barbarus* and *A. tigrinum*, yank for the net GRF and vertical component of the GRF (Fig. 5a – b) approximated sine waves that started close to zero at the beginning of stance, increased to positive yank values before decreasing to an inflection point around 60% of stance, and then decreased to negative yank values before returning to the zero baseline. In contrast, net GRF and vertical component of the GRF for *P. waltl* reached an inflection point at ~50% of stance, and then approached zero again at ~80% of stance, at a time when *A. tigrinum* and *P. barbarus* were both reaching their minimum yank values (i.e., peak in negative yank). The anteroposterior component was comparable between the two salamander forelimbs (~0.006 - 0.008 BW for each percent of stance), which increased faster than the pectoral fin of *P. barbarus* (0.004 ± 0.001 BW for each percent of stance). These findings demonstrate that rates of GRF production in the pectoral appendages of the semi-aquatic salamander, *P. waltl*, were generally similar in magnitude but shifted at different times during stance compared to the other two species.

**Figure 5.**
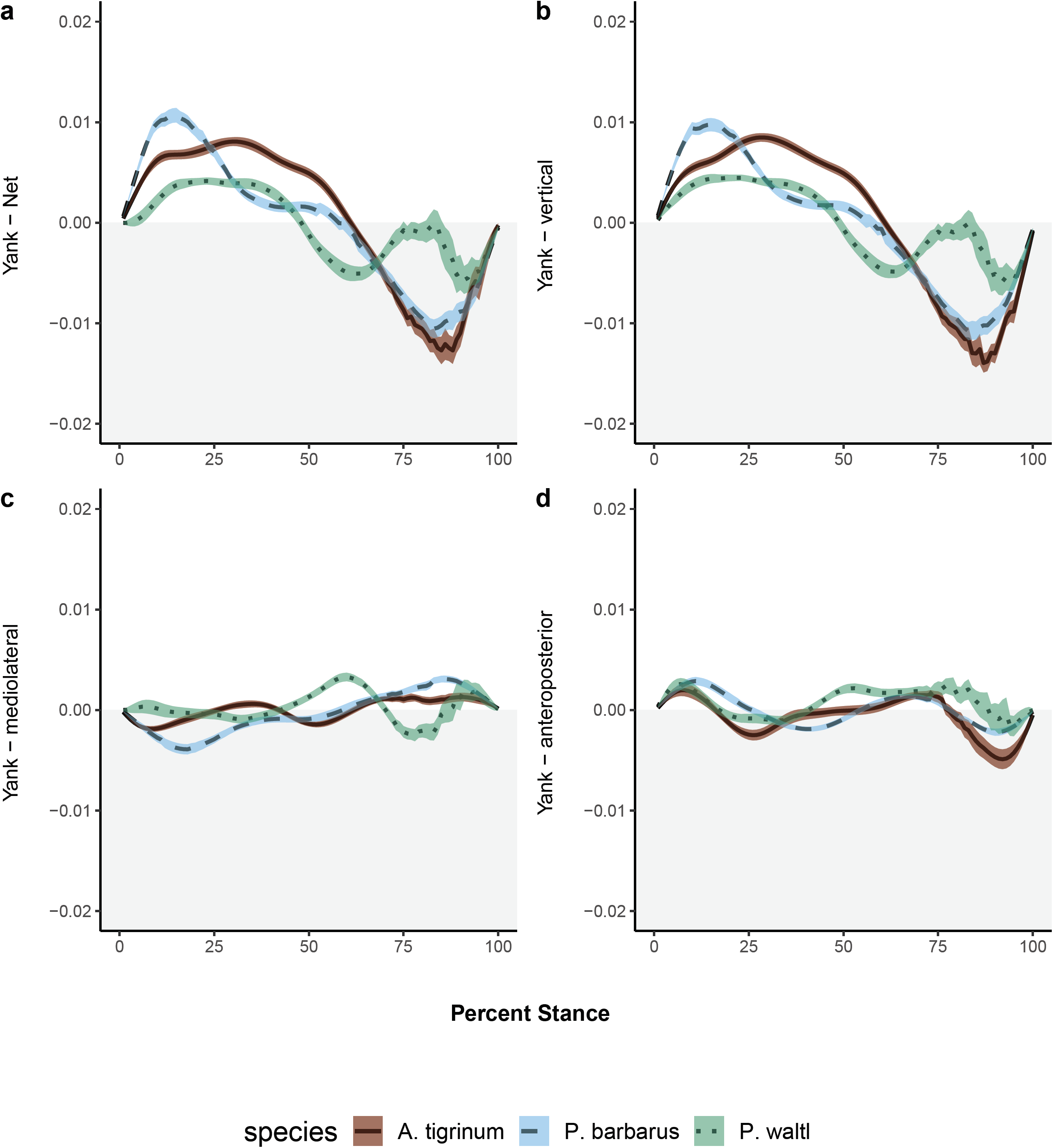
Profiles of the rate of GRF production (‘yank’) in the pectoral appendages during stance. Individual trials were pooled within each species (N = 50 trials for each) and profiles are plotted as the mean ± s.e.m. for each percent of stance. Mediolateral components were set relative to vertical (0°), so negative values indicate a medial direction of the GRF. Anteroposterior components were set relative to vertical (0°), so negative values represent a posterior direction of the GRF. Yank values for *P. waltl* (dotted green lines) tended to span a narrower range of values and did not generally follow the same shape dynamics as the semi-aquatic fish *P. barbarus* (dashed blue lines) and the primarily terrestrial salamander *A. tigrinum* (solid brown lines), which tended to yield comparable data.

**Table 3.** Range of average yank values compared between appendages and species

|  | Vertical |  | Mediolateral |  | Anteroposterior |  | Net |  |
| --- | --- | --- | --- | --- | --- | --- | --- | --- |
| | FE ± s.e.m. | CI | FE ± s.e.m. | CI | FE ± s.e.m. | CI | FE ± s.e.m. | CI |
| **Pectoral - Max** |  |  |  |  |  |  |  |  |
| *P. barbarus* | 0.010 ± 0.001 | 0.008, 0.012 | 0.004 ± 0.001 | 0.003, 0.006 | 0.004 ± 0.001 | 0.002, 0.005 | 0.011 ± 0.001 | 0.008, 0.013 |
| *P. waltl* | 0.009 ± 0.001 | 0.007, 0.011 | 0.006 ± 0.001 | 0.004, 0.008 | 0.008 ± 0.001 | 0.007, 0.010 | 0.009 ± 0.001 | 0.007, 0.012 |
| *A. tigrinum* | 0.012 ± 0.001 | 0.010, 0.014 | 0.004 ± 0.001 | 0.002, 0.005 | 0.006 ± 0.001 | 0.004, 0.008 | 0.012 ± 0.001 | 0.010, 0.014 |
| **Pectoral - Min** |  |  |  |  |  |  |  |  |
| *P. barbarus* | -0.012 ± 0.001 | -0.014, -0.010 | -0.005 ± 0.001 | -0.006, -0.003 | -0.004 ± 0.001 | -0.005, -0.003 | -0.013 ± 0.001 | -0.015, -0.011 |
| *P. waltl* | -0.009 ± 0.001 | -0.011, -0.007 | -0.006 ± 0.001 | -0.008, -0.005 | -0.005 ± 0.001 | -0.006, -0.003 | -0.010 ± 0.001 | -0.012, -0.007 |
| *A. tigrinum* | -0.013 ± 0.001 | -0.016, -0.011 | -0.004 ± 0.001 | -0.006, -0.003 | -0.007 ± 0.001 | -0.008, -0.005 | -0.014 ± 0.001 | -0.016, -0.011 |
| **Pelvic - Max** |  |  |  |  |  |  |  |  |
| *P. waltl* | 0.005 ± 0.001 | 0.002, 0.008 | 0.006 ± 0.001 | 0.005, 0.008 | 0.006 ± 0.001 | 0.004, 0.008 | 0.006 ± 0.001 | 0.003, 0.009 |
| *A. tigrinum* | 0.010 ± 0.001 | 0.007, 0.013 | 0.006 ± 0.001 | 0.005, 0.008 | 0.009 ± 0.001 | 0.007, 0.011 | 0.010 ± 0.001 | 0.007, 0.014 |
| **Pelvic - Min** |  |  |  |  |  |  |  |  |
| *P. waltl* | -0.009 ± 0.001 | -0.011, -0.007 | -0.004 ± 0.0004 | -0.005, -0.003 | -0.006 ± 0.001 | -0.007, -0.005 | -0.009 ± 0.001 | -0.011, -0.007 |
| *A. tigrinum* | -0.014 ± 0.001 | -0.016, -0.012 | -0.004 ± 0.0004 | -0.005, -0.003 | -0.007 ± 0.001 | -0.008, -0.006 | -0.015 ± 0.001 | -0.016, -0.013 |

Sample sizes for the pectoral yank LMM: *P. barbarus* (n = 50), *P. waltl* (n = 50), and *A. tigrinum* (n = 50). Sample sizes for the pelvic yank LMM: *P. waltl* (n = 49), and *A. tigrinum* (n = 49). FE = estimates of the coefficients for the fixed effects based on estimated marginal means with the emmeans::emmeans() function. Statistical analyses were based on the model: lme4::lmer(y ~ species + (1 | individual), REML = True). The confidence interval (CI) is listed as the lower and upper bounds, respectively, that were reported from the LMMs.

Similarities between *A. tigrinum* and *P. waltl* were more evident in the shapes of the yank curves for the hind limbs (Fig. 6). The curves for the net GRF (Fig. 6a) and the vertical component of the GRF (Fig. 6b) approximated sine waves that reached an inflection point at around 25% of stance. The peak for positive yank appears almost 2× higher in *A. tigrinum* compared to *P. waltl* (Fig. 6, Table 3), but the confidence intervals overlapped for the fixed effects. The minima for negative yank in the vertical component and net GRF were shallower in *P. waltl* compared to *A. tigrinum*. As a result, rates of change in GRF production for the hind limb were slower for the semi-aquatic *P. waltl* salamander compared to the terrestrial *A. tigrinum* salamander for the vertical component of the GRF and net GRF, but were generally similar otherwise.

**Figure 6.**
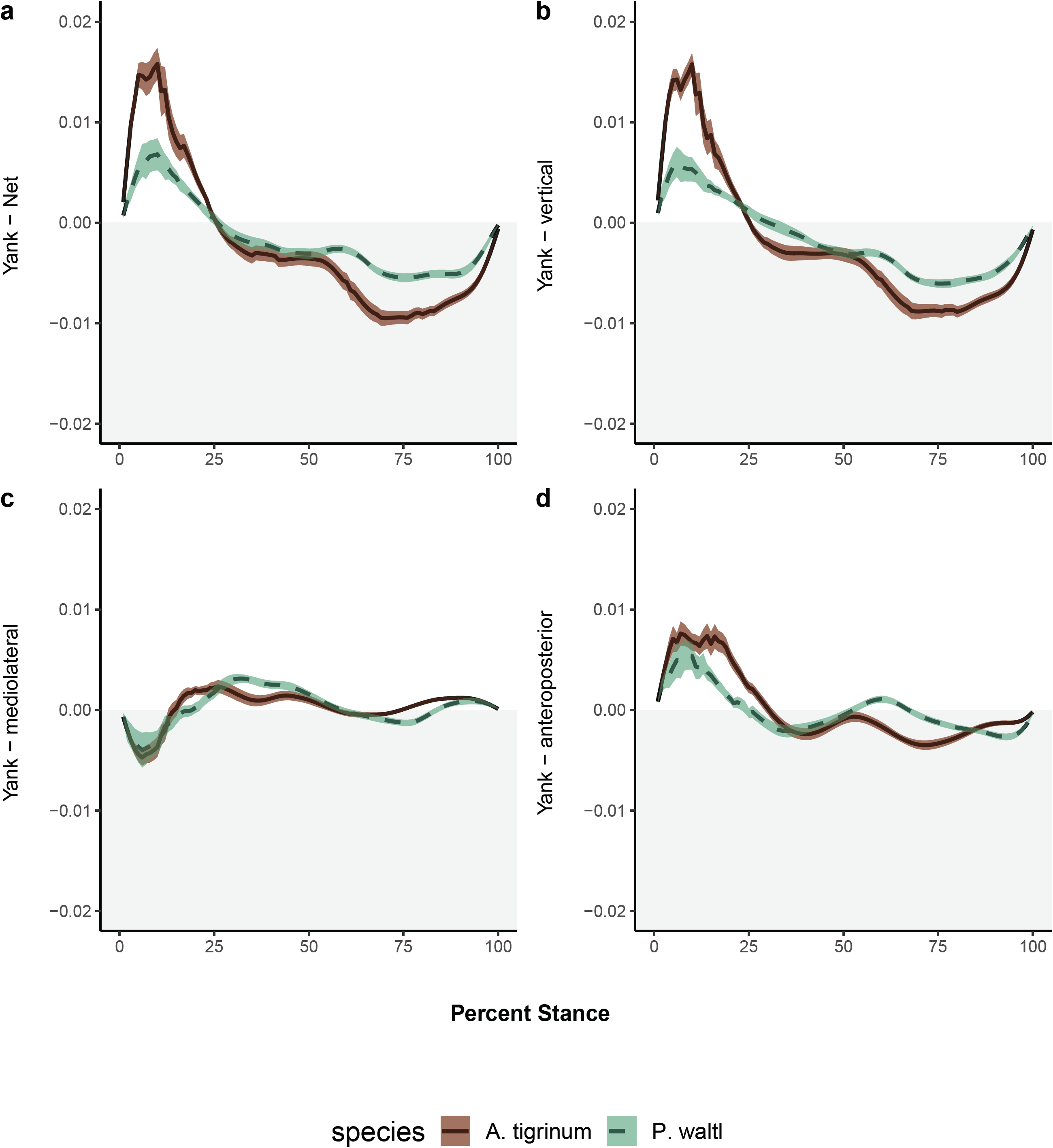
Profiles of the rate of GRF production (‘yank’) in the pelvic appendages during stance. Individual trials were pooled within each species (N = 49 trials for each) and profiles are plotted as the mean ± s.e.m. for each percent of stance. Mediolateral components were set relative to vertical (0°), so negative values indicate a medial direction of the GRF. Anteroposterior components were set relative to vertical (0°), so negative values represent a posterior direction of the GRF. Yank values for *P. waltl* hind limbs (dashed green lines) tended to span a narrower range of values for the vertical component and net GRF, but were otherwise relatively comparable to those from the primarily terrestrial salamander *A. tigrinum* (solid brown lines).

## DISCUSSION

### Distinctions in limb mechanics for semi-aquatic salamanders

*Pleurodeles waltl* is a semi-aquatic salamander that spends most of its life history underwater. Similar to terrestrial salamanders, the predominant acceleratory forces in this semi-aquatic salamander are produced by the hind limb, signifying rear-wheel drive. The orientation of the GRF in the hind limb was about 17° in the anterior direction for both salamanders, but was more medial in *P. waltl* (~17°) compared to *A. tigrinum* (~9°). Similarly, the orientation of the GRF for the pectoral appendages was more medial in the semi-aquatic *P. waltl and P. barbarus* compared to the primarily terrestrial *A. tigrinum* (Table 1). These differences in orientation may affect how loads are applied to the appendages, potentially resulting in the lower vertical component and net magnitude of the GRF in *P. waltl* forelimbs and hind limbs.

Although the forelimbs of both salamanders had a net deceleratory role as evidenced by the negative value for the anteroposterior component of the GRF, this value was larger in magnitude throughout almost all of stance in *P. waltl*, whereas *A. tigrinum* exhibited two periods of slightly net positive values (Fig. 1, Table 1). Comparisons between the limbs of *P. waltl* demonstrate that the hind limbs are the primary propulsors, whereas the forelimbs may be operating as ‘struts’ that act as pivot points to rotate the body. This suggests that semi-aquatic and terrestrial salamanders may be using different mechanisms for propulsion between the forelimbs and hind limbs.

McElroy and colleagues (2014) found that differential limb function occurred in diverse ways across sprawling quadrupeds (e.g., diverse lizard taxa), and that ‘rear-wheel drive’ was one of the only unifying themes among the seven species compared. Instead, differences in the locomotor biomechanics of limbs in running quadrupedal lizards were based on interspecific differences in limb morphology, with a more medial orientation of the GRFs produced by the hind limb in lizard species that had disproportionately longer hind limbs than the forelimbs (McElroy et al., 2014). Differential limb function in sprawling quadrupeds is proposed to increase maneuverability while maintaining stability (McElroy et al., 2014). Although this idea has been proposed in the context of running lizards that may experience instability during rapid turns, similar arguments apply to the locomotion of crabs, enabling them to avoid over-turning moments from hydrodynamic forces (Chen et al., 2006). Future studies could factor environmental variation into comparative analyses of differential limb function to examine how it might evolve in the context of selection pressures presented by different environments.

Disparities in function between fore- and hind appendages, as well as other differences between biomechanical profiles for aquatic and terrestrial species, could relate to the different demands imposed by the primary environments in which the appendages of these taxa function. For example, point estimates for the medial orientation of the peak GRF in semi-aquatic *P. waltl* limbs (14-16°) falls between that of *P. barbarus* fins (17°) and most previously evaluated tetrapod limbs (<11°), including primarily terrestrial *A. tigrinum* salamanders (Fig. 7, Tables 1 - 2). A shift to a GRF directed less medially could reduce joint moments and, thus, the stresses experienced by the appendicular bones during terrestrial locomotion (Kawano and Blob, 2013). However, the greater medial orientation of the GRF in semi-aquatic *P. waltl* likely relates to the greater lateral spread of their distal limb segments compared to terrestrial taxa, so that the feet are placed lateral to the elbow or knee joint during stance (Fig. S1A), rather than directly below these joints (as in terrestrial salamanders: Fig. 1A, B in Kawano and Blob, 2013). A recent comparison of terrestrial locomotion in salamanders found that walking mechanics and muscle activation patterns were generally similar among species, but that some interspecific differences emerged (Pierce et al., 2020). For instance, maximum extensions of the wrist (160°) and ankle (159°) were higher in *P. waltl* than *A. tigrinum* (150° and 150°, respectively) despite its elbow and knee having greater (55° vs. 30°) or equal (55°) total maximum extensions, respectively (Pierce et al., 2020). Given that this more pronounced sprawling limb posture is also found in the mudskipper fish, a more medial orientation of the GRF also might be found in other semi-aquatic taxa. For example, Ashley-Ross and Bechtel (Ashley-Ross and Bechtel, 2004) found that *Taricha torosa* newts also had greater lateral spread of the limbs, caused by more flattened angles of the limb elements, when walking in water than on land.

**Figure 7.**
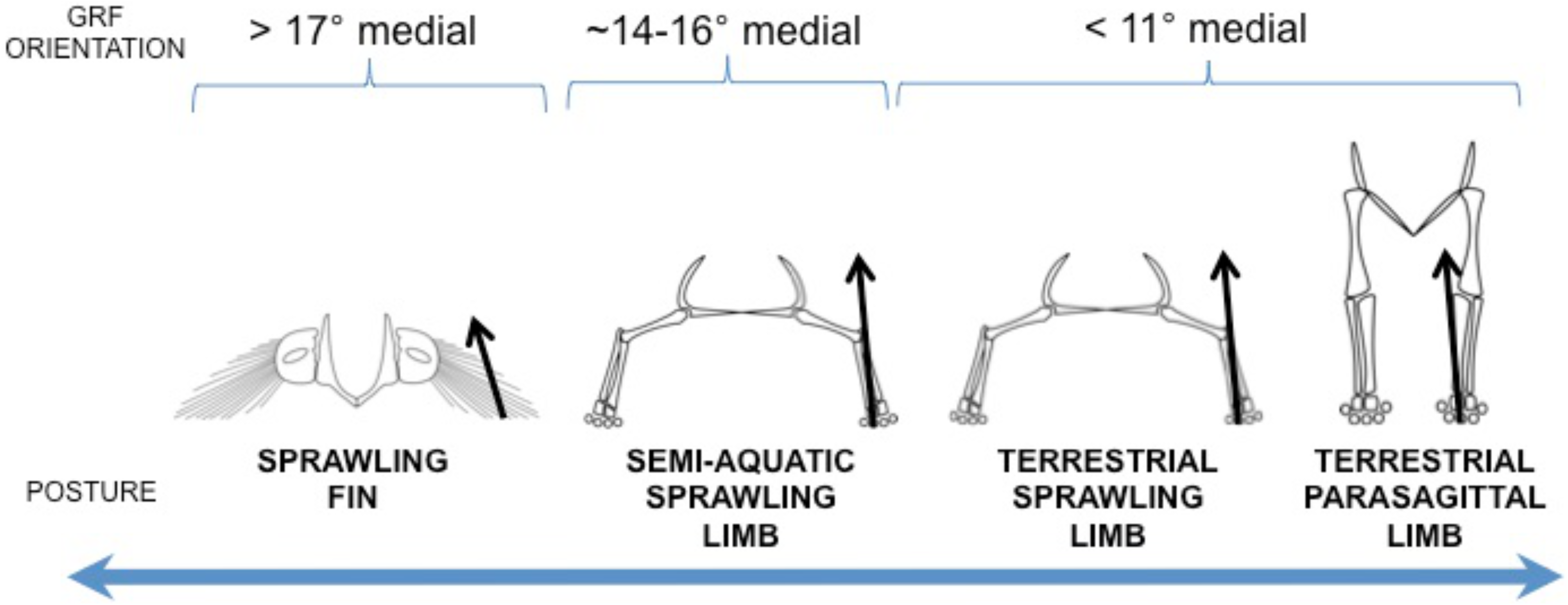
Data on GRF production in semi-aquatic *P. waltl* salamanders contribute towards a broader perspective on the evolution of the locomotor biomechanics across the fish-tetrapod transition. The orientation of the GRF (represented as a black arrow) in the medial direction was historically thought to be evolutionarily conserved across tetrapods, despite variation in limb postures. It was unclear whether the same pattern was found in fishes, but our previous work found that GRFs produced by pectoral fins in mudskipper fishes were more medial, likely resulting from their more sprawled posture compared to tetrapods. GRF data from *P. waltl* in this study demonstrate that the more medial orientation of GRFs is not exclusive to fishes, suggesting that ecology may have an important role in shaping the kinetics during terrestrial locomotion.

The observation that *P. waltl* maintains a more extended sprawling posture on land may mean that some species have greater stereotypy (Wainwright et al., 2008) in their locomotor biomechanics compared to others, with the degree of flexibility influenced by how often habitat transitions occur during a season. The broadening of the gait pattern that would result from such lateral foot placement might convey additional stability against currents or other flows in aquatic habitats (Martinez et al., 1998) by reducing pitching and rolling (Chen et al., 2006). However, when on land, species that tend to be more aquatic may not be able to adjust to the more upright orientations of distal limb segments that are seen in terrestrial taxa (Kawano and Blob, 2013). Producing more acute limb angles could facilitate elevating the body off the ground, and shift the bone loading regime to reduce bending and increase compression (Ashley-Ross and Bechtel, 2004; Kawano and Blob, 2013). Lateral spread of the distal appendage may also contribute to the high medial orientation of the GRF in mudskippers (Fig. 1C in Kawano and Blob, 2013), but furthermore could prevent rolling of the body axis during terrestrial crutching, given the lack of extended posterior appendages in this taxon. Thus, shifts to less sprawling limb posture could have major biomechanical consequences that could facilitate terrestrial locomotion.

Similarities in the overall shapes of the GRF and yank produced by individual hind limbs suggest that rates of GRF production are consistent across hind limbs of semi-aquatic and terrestrial salamander species, but the different peaks observed in these curves indicate that the maxima differ between them. The similarities in force production between semi-aquatic and terrestrial hind limbs could indicate that the use of the hind limbs as a primary propulsor imposes strong selection on kinetics for acceleration, but that other aspects, such as maneuverability and body support, can be altered by changing the GRFs and yank. For example, the smaller absolute value for negative yank of the vertical component of the GRF and net GRF for the hind limbs in *P. waltl*, compared to *A. tigrinum*, suggest that there may be steadier loading regimes in the former. Loading the appendages in more predictable ways can have numerous advantages (Granatosky et al., 2020). For instance, McNeil Alexander proposed that higher safety factors (margins of protection against failure) may be found in structures that experience more variable loading regimes (Alexander, 1997). Consequently, steadier loading regimes could reduce the safety factors needed in the limb bones and, therefore, reduce the energetic costs to produce the higher safety factors. Such factors might be advantageous if loads were elevated due to the highly sprawling posture of *P. waltl* (Fig. 7). The limb bones of *A. tigrinum* have been found to possess generally high safety factors against failure (Blob et al., 2014; Kawano et al., 2016; Sheffield and Blob, 2011) and studies are ongoing to determine whether a similar pattern is found in semi-aquatic salamanders.

The rate of load application can also affect the load magnitudes experienced by bones. Strain magnitude is positively correlated with strain rate for both cyclic and discrete loading regimes in vertebrates (Aiello et al., 2015), so smaller yank values in *P. waltl* should be associated with slower strain rates and lower peak strain magnitudes. Lower peak strains would keep bones further from failure, whereas slower strain rates could affect bone mass (Rubin and Lanyon, 1982). Thus, peak strains could have a direct impact on the safety factors of the limb bones, and strain rates could affect the remodelling of bones in response to different loading regimes. Strain rate has been found to be a main factor driving bone remodelling and was positively correlated with higher Young’s modulus values (Aiello et al., 2015), which would confer greater stiffness and greater resistance to bending in bones. Such properties would be advantageous for terrestrial taxa that must habitually withstand the downward effects of gravity on land. Semi-aquatic taxa that spend most of their time underwater might not need higher stiffnesses in their limb bones while their bodies are primarily being supported by buoyancy; however, such traits could be beneficial during the terrestrial eft phase of many newt species (including *P. waltl*). Future studies that examine changes in the mechanical properties and loading regimes of limb bones across ontogenetic stages within salamanders and between species with different life histories (e.g., direct vs. indirect developers) would provide further insights into the factors that shape limb function.

#### Paleontological implications

Evidence from the fossil record suggests that terrestrial adaptations first appeared in the anterior regions of the body (Lebedev, 1997), but how rear-wheel drive evolved within stem tetrapods is still debated. Anatomical evaluations of some of the earliest stem tetrapods, such as the elpistostegalid fish *Panderichthys* (Boisvert, 2005) and the Devonian tetrapod *Ichthyostega* (Pierce et al., 2012), indicate that the pelvic appendages were likely not effective propulsors on land. As a result, front-wheel drive has been proposed in the early stages of terrestrial locomotion for at least some early tetrapods (Boisvert, 2005; Nyakatura et al., 2014). However, empirical work on the African lungfish (*Protopterus annectens*) suggests that rear-wheel drive likely evolved for aquatic locomotion when vertebrates were still fully aquatic (King et al., 2011). Further, recent paleontological examinations of the pelvic girdle of the elpistostegalid tetrapodomorph fish *Tiktaalik* (a relative of *Panderichthys*) indicate that it exhibited a mosaic of tetrapod-like and fish-like characteristics, including precursors for achieving rear-wheel drive (Shubin et al., 2014). Given the morphological diversity across tetrapodomorphs, it is appropriate to evaluate a range of extant taxa as functional models that represent different locomotor strategies. Our GRF data from semi-aquatic *P. waltl* limbs build upon previous work on the kinetics of mudskipper pectoral fins and terrestrial salamander limbs (Kawano and Blob, 2013) to offer additional insights for interpreting evolutionary patterns in the incipient stages of terrestrial locomotion, providing a functional model for semi-aquatic tetrapods that exhibit locomotor biomechanics intermediate between those of finned taxa and crownward tetrapods.

Taxa with intermediate morphologies and locomotor biomechanics may be particularly relevant for stem tetrapods. Morphological comparisons across the humeri of 40 tetrapodomorphs spanning the water-to-land transition demonstrated that stem tetrapods had intermediate morphologies that placed them in a valley between two adaptive peaks (one for aquatic fishes and second for terrestrial crown tetrapods) (Dickson et al., 2021). This was a likely transitional state before natural selection was strong enough to favor the movement of stem tetrapods to the adaptive peak representing the terrestrial characteristics now associated with crown tetrapods (Dickson et al., 2021). Work by Nyakatura and colleagues (Nyakatura et al., 2014) suggests that tetrapods may have gone through an intermediate stage during the transition from front-wheel drive to rear-wheel drive. Specifically, their work evaluated the limb mechanics of a sprawling, belly-dragging lizard, and proposed that belly dragging could have allowed early tetrapods to move on land using appendicular muscles that were less developed (Nyakatura et al., 2014). This intermediate belly-dragging stage of front-wheel drive among early tetrapods would have allowed initial capacities for terrestrial locomotion, after which the role of rear-wheel drive gradually increased. Our findings from the semi-aquatic salamander, *P. waltl*, may provide a model for a subsequent stage after belly-dragging with front-wheel drive, in which rear-wheel drive has been adopted on land but the forelimbs have not yet acquired fully terrestrial limb mechanics.

Kinetic data from the semi-aquatic salamander may serve as a foundation for following up with two hypotheses regarding how terrestrial locomotion evolved (discussed in Pierce et al., 2013). The first hypothesis suggested that the first locomotor mode on land was a trot with lateral bending of the axial system, producing a traveling wave and limbs that acted as ‘struts’. The second hypothesis proposed a lateral sequence walk involving a standing wave, with the limbs generating propulsion. Given that the semi-aquatic forelimb was deceleratory while the hind limb was acceleratory (Tables 1-2, Figs. 1 - 4), *P. waltl* may be using a modified standing wave in which the hind limbs are generating forward propulsion while the forelimbs are being used as ‘struts’. Such disparity in the propulsive roles of the limbs is not as pronounced in the terrestrial salamander (Table 1 in Kawano and Blob, 2013) and may be due to the different habitats and ecologies that are associated with these species. Similarly, Pierce and colleagues (2020) found interspecific differences in both kinematic and muscle activation patterns during terrestrial walking and proposed that the degree of terrestriality, or other factors related to ecology, may be driving such variation. A gait that was similar to the one used by *P. waltl* may have allowed the earliest limbed tetrapods to traverse the terrestrial environment with a musculoskeletal system that still primarily functioned for underwater behaviors (McInroe et al., 2016), potentially representing an intermediate stage between sarcopytergian fishes that could accomplish rear-wheel drive underwater (King et al., 2011) and crownward tetrapods that used rear-wheel drive on land.

How functional changes evolve has been considered in a variety of systems. Historically, the evolution of locomotor posture has been viewed as a sequential series of gradual evolutionary steps, leading from sprawling to upright (Charig, 1972). However, more recent work highlighted the potential for intermediate taxa to exhibit a broader range of capabilities between the ends of this functional continuum, rather than a graded series of incremental changes between them (Blob, 2001; Kemp, 1978). Hind limb function in the tetrapodomorph fish *Tiktaalik* has been described with a wide range of capacities that may have been possible due to the relaxed constraints placed upon musculoskeletal systems to overcome gravitational loads (Shubin et al., 2014), potentially indicating intermediate flexibility in locomotor function for an early stage of the fin-to-limb transition that allowed for greater rotations about the hip than tetrapods (and greater stability in the shoulder compared to the hip), but reduced ability to withstand locomotor stresses. Such disparity in limb function is reflected in our data from *P. waltl*, which suggest that the forelimbs and hind limbs have different functional roles for generating forward movement. Moreover, these changes may not have been strictly coupled to evolutionary changes in appendicular structure since such stark differences between the forelimbs and hind limbs were not found in the GRFs for terrestrial *A. tigrinum* salamanders. Synthesis of empirical data from biomechanical experiments and morphological analyses of fossils from palaeontology, therefore, holds promise for developing more comprehensive reconstructions of the transformations in the vertebrate musculoskeletal system that led to digit-bearing tetrapods becoming terrestrial (Nyakatura et al., 2019) and the causes and effects of functional evolution more broadly.

## CONCLUSION

Our results provide novel insights into the functional changes associated with tetrapods becoming terrestrial, with broader implications for whether functional capabilities could be closely associated with major structural changes (e.g., transition from fin to limb) or could occur through gradual changes that included intermediate stages. The kinetics of terrestrial walking in semi-aquatic *P. waltl* salamanders differed between the limbs, with the hind limbs sharing numerous similarities to the hind limbs of primarily terrestrial salamanders but the forelimbs exhibiting kinetics that were distinct from the pectoral appendages of semi-aquatic fish and primarily terrestrial salamanders. These data provide additional context to account for differential limb function in models of musculoskeletal function, including those that use salamanders as a blueprint for estimating the locomotor function of early tetrapods. Together, our results present new context for the functional consequences of different locomotor strategies that are possible in functional analogs of tetrapodomorphs that spanned the water-to-land transition.

## ACKNOWLEDGMENTS

We thank Zach Marion, Jim Fordyce, and Yu Xia for helpful discussions on statistical analyses that were considered for this study. We are also grateful to Margaret Ptacek, Miriam Ashley-Ross, and anonymous reviewers for feedback on earlier drafts of this manuscript. An earlier draft of this manuscript was submitted by S.M.K. in partial fulfilment of a doctoral dissertation at Clemson University.

## DATA ACCESSIBILITY

All R code are open access via GitHub (https://github.com/MorphoFun/kraken), with the specific scripts for this manuscript available in the “compareGRFs” folder. The raw files for the original force data were too large to include on GitHub but will be made available on Open Science Framework (https://osf.io/) upon publication of the results.

## AUTHOR CONTRIBUTIONS

S.M.K and R.W.B. designed the experiment. S.M.K. ran the experiments, wrote the R scripts, and analyzed the data. S.M.K. and R.W.B. contributed to interpreting the results, writing the manuscript, and approving all drafts of the manuscript.

